# Gene model for the ortholog of *tgo* in *Drosophila miranda*

**DOI:** 10.1101/2025.08.04.668519

**Authors:** Graham M. Jones, Abbie A. Giunta, Hanah Mastandrea, Jacqueline K. Wittke-Thompson

## Abstract

Gene model for the ortholog of *tango* (*tgo*) in the April 2013 (UC Berkeley DroMir_2.2/DmirGB2) Genome Assembly (GenBank Accession: GCA_000269505.2) of *Drosophila miranda*. This ortholog was characterized as part of a developing dataset to study the evolution of the Insulin/insulin-like growth factor signaling pathway (IIS) across the genus *Drosophila* using the Genomics Education Partnership gene annotation protocol for Course-based Undergraduate Research Experiences.

## Introduction

*This article reports a predicted gene model generated by undergraduate work using a structured gene model annotation protocol defined by the Genomics Education Partnership (GEP; *thegep.org*) for Course-based Undergraduate Research Experience (CURE). The following information in quotes may be repeated in other articles submitted by participants using the same GEP CURE protocol for annotating Drosophila species orthologs of Drosophila melanogaster genes in the insulin signaling pathway*.

“In this GEP CURE protocol students use web-based tools to manually annotate genes in non-model *Drosophila* species based on orthology to genes in the well-annotated model organism fruit fly *Drosophila melanogaster*. The GEP uses web-based tools to allow undergraduates to participate in course-based research by generating manual annotations of genes in non-model species (Rele et al., 2023). Computational-based gene predictions in any organism are often improved by careful manual annotation and curation, allowing for more accurate analyses of gene and genome evolution (Mudge and Harrow 2016; Tello-Ruiz et al., 2019). These models of orthologous genes across species, such as the one presented here, then provide a reliable basis for further evolutionary genomic analyses when made available to the scientific community.” (Myers et al., 2024).

“The particular gene ortholog described here was characterized as part of a developing dataset to study the evolution of the Insulin/insulin-like growth factor signaling pathway (IIS) across the genus *Drosophila*. The Insulin/insulin-like growth factor signaling pathway (IIS) is a highly conserved signaling pathway in animals and is central to mediating organismal responses to nutrients (Hietakangas and Cohen 2009; Grewal 2009).” (Myers et al., 2024).

“The Drosophila *tango* (CG11987, FBgn0264075) gene encodes a bHLH-PAS protein that controls CNS midline and tracheal development (Sonnenfeld et al., 1997). Both cell culture and in vivo studies have shown that a DNA enhancer element acts as a binding site for both Single-minded∷Tango and Trachealess∷Tango heterodimers and functions in controlling CNS midline and tracheal transcription. Isolation and analysis of tango mutants reveal CNS midline and tracheal defects. In addition, the bHLH-PAS proteins Similar (Sima) and Tango (Tgo) function as HIF-α and HIF-β homologues, respectively, in a conserved hypoxia-inducible transcriptional response in *Drosophila melanogaster* that is homologous to the mammalian HIF-dependent response (Lavista-Llanos et al., 2002). Insulin has been shown to activate HIF-dependent transcription, both in *Drosophila* S2 cells and in living *Drosophila* embryos, and this effect is mediated by PI3K-AKT and TOR pathways. Overexpression of dAKT and dPDK1 in normoxic embryos provoked a major increase in Sima nuclear localization, mimicking the effect of a hypoxic treatment (Dekanty et al., 2005).” (Lawson et al., 2025).

“*D. miranda* (NCBI:txid 7229) is part of the *psuedoobscura* species subgroup within the *obscura* species group in the subgenus *Sophophora* of the genus *Drosophila* (Sturtevant 1942; Buzzati-Traverso and Scossiroli 1955). It was first described by Dobzhansky in 1935. Like other members of the species subgroup, it is endemic to the New World distributed through Canada, the USA, and Mexico, sympatric with its sibling species *D. pseudoobscura* (Markow and O’Grady 2005), living in temperate forest environments.” (Lawson et. al., 2024).

We propose a gene model for the ortholog in *D. miranda* of the *D. melanogaster* tango (*tgo*) gene. The genomic region of the ortholog corresponds to the uncharacterized protein XP_017141376.1 (Locus ID LOC108155207) in the April 2013 (UC Berkeley DroMir_2.2/DmirGB2) Genome Assembly of *D. miranda* (GCA_000269505.2, Alekseyenko et al., 2013; Zhou and Bachtrog 2012). This model is based on RNA-Seq data from *D. miranda* (Alekseyenko et al., 2013; Zhou and Bachtrog 2012; SRP009365) and *tgo* in *D. melanogaster* using FlyBase release FB2024_02 (GCA_000001215.4; Gramates et al., 2022; Jenkins et al., 2022; Larkin et al., 2021).

### Synteny

*tgo* occurs on chromosome 3R in *D. melanogaster* and is flanked by *hyx* and *neur* upstream and *CG11986* and *Sf3b5* downstream. We determined that the putative ortholog of *tgo* is found on scaffold CM001518 in *D. miranda* with LOCUS ID LOC108155207 (via *tblastn* search with an E-value of 0.0 and 95.21% identity), where it is surrounded by LOCUS IDs LOC108155719, LOC108156139, LOC108155210, LOC108155208 which correspond to *hyx, neur, CG11986*, and *Kcmf1* in *D. melanogaster* with percent identities of 90.93%, 88.21%, 81.49%, and 80.96%, E-values of 0.0 for all, and XP IDs of XP_017142211.1, XP_017142942.1, XP_017141383.1, XP_033245776.1, respectively as determined by *blastp* (Figure 1A, Altschul et al., 1990). We believe this is the correct ortholog assignment for *tgo* in *D. miranda* because of the high degree of local synteny, both in gene orientation and relative location. It should be noted that the second most downstream gene in *D. miranda, Kcmf1*, does not appear to be orthologous to the second most downstream gene in *D. melanogaster, Sf3bf*. However, the other genes within the genomic neighborhood of *tgo* in *D. miranda* appear to be orthologous to the genes in the same respective relative location in *D. melanogaster*.

**Figure 1.**
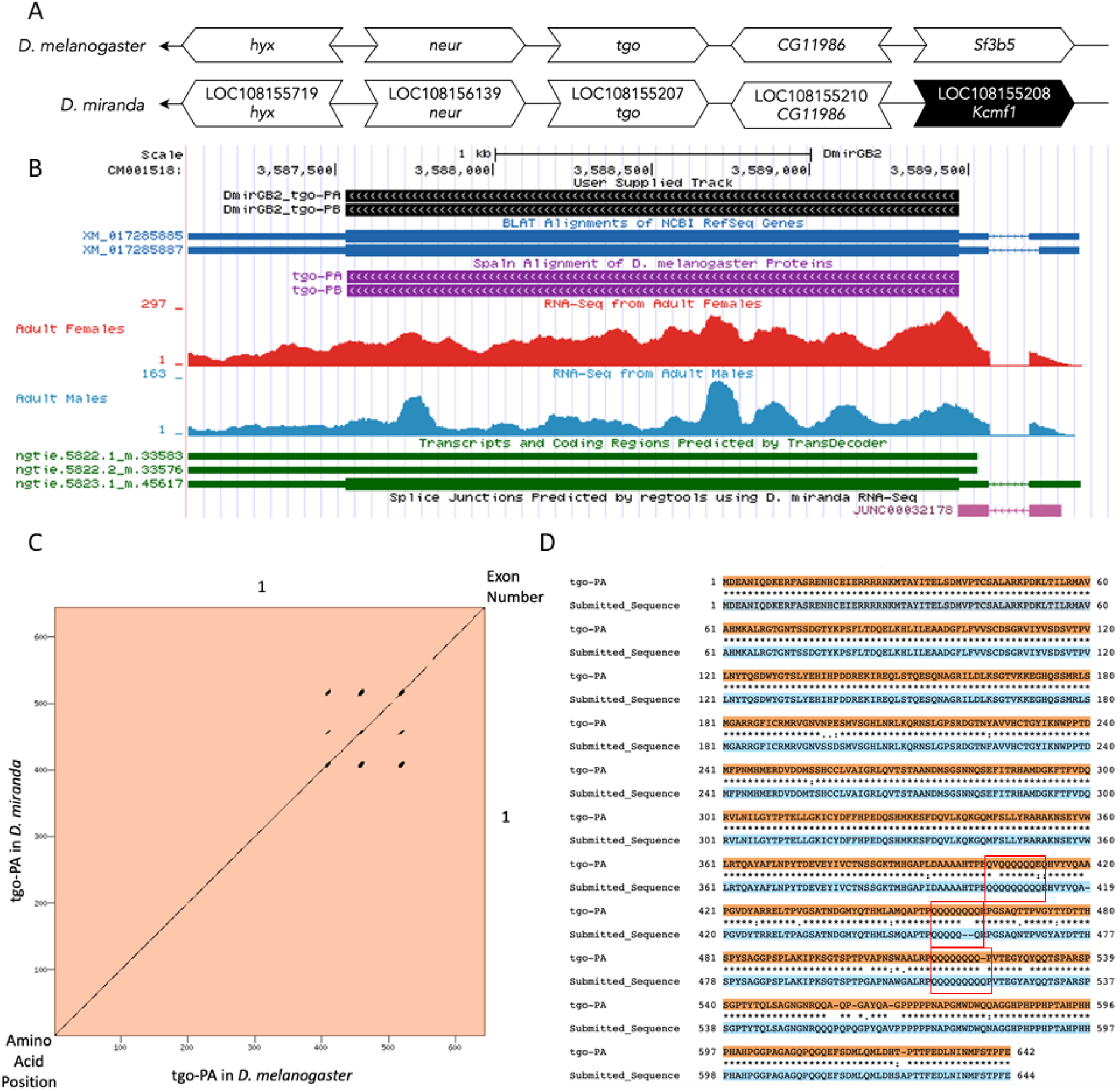
*tgo* gene model comparison between *Drosophila miranda* and *Drosophila melanogaster* orthologs. **(A) Synteny comparison of the genomic neighborhoods for *tgo* in *Drosophila melanogaster* and *D. miranda***. Gene arrows pointing in the same direction as target gene in both *D. miranda* and *D. melanogaster* are on the same strand as the target gene; gene arrows pointing in the opposite direction are on the opposite strand. The thin underlying arrows pointing to the left indicate that *tgo* is on the negative (−) strands. White arrows in *D. miranda* indicate the locus ID and orthology to the corresponding gene in *D. melanogaster*, and black arrows indicate a non-orthologous gene. Gene symbols given in the *D. miranda* gene arrows indicate the orthologous gene in *D. melanogaster*, while the locus identifiers are specific to *D. miranda*. **(B) Gene Model in UCSC Track Hub** (Raney et al. 2014): the gene model in *D. miranda* (black), Spaln of D. melanogaster Proteins (purple, alignment of refseq proteins from *D. melanogaster*), BLAT alignments of NCBI RefSeq Genes (blue, alignment of refseq genes for *D. miranda*), RNA-Seq from adult females and males (red and blue, alignment of Illumina RNA-Seq reads from *D. miranda*), and Transcripts (green) including coding regions predicted by TransDecoder and Splice Junctions Predicted by regtools using *D. miranda* RNA-Seq (Alekseyenko et al., 2013; Zhou and Bachtrog 2012; SRP009365). The splice junction shown has a read-depth of 105, indicated in pink. The custom gene model (User Supplied Track) is indicated in black with exon depicted with wide boxes, intron with narrow lines (arrows indicate direction of transcription). **(C) Dot Plot of tgo-PA in *D. melanogaster* (*x*-axis) vs. the orthologous peptide in *D. miranda* (*y*-axis)**. Amino acid number is indicated along the left and bottom; coding-exon number is indicated along the top and right, and exons are also highlighted with alternating colors. A region of insertion/deletions and single-codon repeats is indicated with parallel line segments. **(D) Protein alignment of tgo-PA in *D. miranda* against tgo-PA in *D. melanogaster***. Identical amino acids indicated with a star (^*^), very similar amino acids indicated with a colon (:), somewhat similar amino acids indicated with a period (.), and insertions or deletions indicated with a dash (−). Repeating glutamine elements indicated with a red box.

### Protein Model

*tgo* in *D. miranda* has two identical protein coding isoforms (tgo-PA and tgo-PB; Figure 1B). Isoforms tgo-PA and tgo-PB contain one CDS. The CDS number and isoform count are conserved relative to the ortholog in *D. melanogaster*. The sequence of tgo-PA in *D. miranda* has 93.51% identity (E-value: 0.0) with the protein coding isoform tgo-PA in *D. melanogaster*, as determined by *blastp* (Figure 1C). A region containing several glutamine repeats was identified towards the C-terminus end of the protein (Figure 1D). This gene model can also be seen within the target genome at this TrackHub.

## Methods

“Detailed methods including algorithms, database versions, and citations for the complete annotation process can be found in Rele et al. (2023). Briefly, students use the GEP instance of the UCSC Genome Browser v.435 (https://gander.wustl.edu; Kent et al., 2002; Navarro Gonzalez et al., 2021) to examine the genomic neighborhood of their reference IIS gene in the *D. melanogaster* genome assembly (Aug. 2014; BDGP Release 6 + ISO1 MT/dm6). Students then retrieve the protein sequence for the *D. melanogaster* reference gene for a given isoform and run it using *tblastn* against their target *Drosophila* species genome assembly on the NCBI BLAST server (https://blast.ncbi.nlm.nih.gov/Blast.cgi; Altschul et al., 1990) to identify potential orthologs. To validate the potential ortholog, students compare the local genomic neighborhood of their potential ortholog with the genomic neighborhood of their reference gene in *D. melanogaster*. This local synteny analysis includes at minimum the two upstream and downstream genes relative to their putative ortholog. They also explore other sets of genomic evidence using multiple alignment tracks in the Genome Browser, including BLAT alignments of RefSeq Genes, Spaln alignment of *D. melanogaster* proteins, multiple gene prediction tracks (e.g., GeMoMa, Geneid, Augustus), and modENCODE RNA-Seq from the target species. Detailed explanation of how these lines of genomic evidenced are leveraged by students in gene model development are described in Rele et al. (2023). Genomic structure information (e.g., CDSs, intron-exon number and boundaries, number of isoforms) for the *D. melanogaster* reference gene is retrieved through the Gene Record Finder (https://gander.wustl.edu/~wilson/dmelgenerecord/index.html; Rele et al., 2023). Approximate splice sites within the target gene are determined using *tblastn* using the CDSs from the *D. melanogaste*r reference gene. Coordinates of CDSs are then refined by examining aligned modENCODE RNA-Seq data, and by applying paradigms of molecular biology such as identifying canonical splice site sequences and ensuring the maintenance of an open reading frame across hypothesized splice sites. Students then confirm the biological validity of their target gene model using the Gene Model Checker (https://gander.wustl.edu/~wilson/dmelgenerecord/index.html; Rele et al., 2023), which compares the structure and translated sequence from their hypothesized target gene model against the *D. melanogaster* reference gene model. At least two independent models for a gene are generated by students under mentorship of their faculty course instructors. Those models are then reconciled by a third independent researcher mentored by the project leaders to produce the final model. Note: comparison of 5’ and 3’ UTR sequence information is not included in this GEP CURE protocol.” (Gruys et al., 2025)

### Supplemental Files

1. Zip file containing a FASTA, PEP, GFF files for the gene model
2. Figure 1 in high resolution

### Metadata

Bioinformatics, Genomics, *Drosophila*, Genotype Data, New Finding

## Supporting information

Gene model data files

## Acknowledgements

We would like to thank Wilson Leung for developing and maintaining the technological infrastructure that was used to create this gene model and Laura K. Reed for overseeing the project. Thank you to [FlyBase](https://flybase.org/) for providing the definitive database for *Drosophila melanogaster* gene models. Further, we would like to thank the editors and developers at the journal *microPublication: Biology* for assistance in developing the template for these single gene ortholog publications.

## Funding

This material is based upon work supported by the National Science Foundation (1915544) and the National Institute of General Medical Sciences of the National Institutes of Health (R25GM130517) to the Genomics Education Partnership (GEP; https://thegep.org/; PI-LKR). Any opinions, findings, and conclusions or recommendations expressed in this material are solely those of the author(s) and do not necessarily reflect the official views of the National Science Foundation nor the National Institutes of Health.

## Notes

https://gander.wustl.edu/cgi-bin/hgTracks?db=DmirGB2&lastVirtModeType=default&lastVirtModeExtraState=&virtModeType=default&virtMode=0&nonVirtPosition=&position=CM001518%3A3587036%2D3589970&hgsid=16717826_PNfDELq625JM7GIuOJeJjkvQjqGr

### Competing Interest Statement

The authors have declared no competing interest.

### Summary of Updates

Updated title and abstract with minor changes to be consistent.

